# True Colors: commercially-acquired morphological genotypes reveal hidden allele variation among dog breeds, informing both trait ancestry and breed potential

**DOI:** 10.1101/654343

**Authors:** Dayna L Dreger, Blair N Hooser, Angela M Hughes, Balasubramanian Ganesan, Jonas Donner, Heidi Anderson, Lauren Holtvoigt, Kari J Ekenstedt

## Abstract

Direct-to-consumer canine genetic testing is becoming increasingly popular among dog owners. The data collected therein provides intriguing insight into the current status of morphological variation present within purebred populations. Mars WISDOM PANEL^TM^ data from 11,790 anonymized dogs, representing 212 breeds and 4 wild canine species, were evaluated at genes associated with 7 coat color traits and 5 physical characteristics. Frequencies for all tested alleles at these 12 genes were determined by breed and by phylogenetic grouping. A sub-set of the data, consisting of 30 breeds, was divided into separate same-breed populations based on country of collection, body size, coat variation, or lineages selected for working or conformation traits. Significantly different (p ≤ 0.00167) allele frequencies were observed between populations for at least one of the tested genes in 26 of the 30 breeds. Next, standard breed descriptions from major American and international registries were used to determine colors and tail lengths (e.g. genetic bobtail) accepted within each breed. Alleles capable of producing traits incongruous with breed descriptions were observed in 143 breeds, such that random mating within breeds has probabilities of between 4.9e^−7^ and 0.25 of creating undesirable phenotypes. Finally, the presence of rare alleles within breeds, such as those for the recessive black coloration and natural bobtail, was combined with previously published identity-by-decent haplotype sharing levels to propose pathways by which the alleles may have spread throughout dog breeds. Taken together, this work demonstrates that: 1) the occurrence of low frequency alleles within breeds can reveal the influence of regional or functional selection practices; 2) it is possible to trace the mode by which characteristics have spread across breeds during historical breed formation; and 3) the necessity of addressing conflicting ideals in breed descriptions relative to actual genetic potential is crucial.

**Author Summary:** From the sleek Doberman Pinscher to the coifed Poodle, the sunny Golden Retriever to the aristocratic Pekingese, the world of purebred dogs offers options that appeal to nearly all aesthetics. Pure dog breeds, of which there are over 400 worldwide, are created through selective breeding over multiple generations, toward an ideal goal of temperament, behavior, and physical appearance. Written descriptions of these breed-specific ideals are produced and maintained by dedicated breed enthusiasts, and provide guidelines that direct breeders in their selection choices. However, the genetic mechanisms that produce the spectrum of canine colors and patterns, sculpt the small triangular ears of a Siberian Husky or the long soft ears of a Basset Hound, are complicated and intertwined. This means that some breeds can carry rare, hidden traits for many generations, hampering selection efforts toward uniformity. We have determined the genotypes of >11,000 dogs, representing >200 breeds, at 12 genes that impact coat color and physical characteristics. In doing so, we can now present realistic trait frequencies within each breed, report occurrences of gene variants that can produce undesirable traits, and draw conclusions about the historic spread of characteristics across modern related breeds and those with distant shared ancestry.

## Introduction

Guided by human selection, the domestic dog (*Canis lupus familiaris*) has become one of the most physically diverse species, with hundreds of recognized breeds, differentiated from each other by specific morphological and behavioral characteristics (1). One of the most readily accessible and easily visualized defining characteristics of dog breeds is in their presentation of pigmentation and color patterns. The nomenclature used currently to refer to dog pigmentation genetics was outlined by Little in 1957 (2). Scientific advancement since that time has allowed for the validation of many of his postulated loci and the identification of causal genetic variants for many of the common coat colors in the species. These coat color and trait variants are increasingly included in commercially-available genetic test panels, the emergence of which has contributed greatly to the acceptance and implementation of genetic screening of purebred dogs by owners, breeders, and veterinarians. In addition to providing valuable information regarding the genetic status of potential breeding dogs in terms of disease state and morphological traits, the combined genotype databases collected by these commercial entities provide a valuable resource for monitoring the diversity of selected alleles, such as those driving coat color, within breeds.

The entirety of coat color patterns in mammals consist of spatial and temporal production of phaeomelanin, yellow- to red-based pigment, and eumelanin, black- to brown-based pigment, controlled by the interaction of multiple genes. In dogs, variation at the *Agouti Signaling Protein* (*ASIP*) gene determines the distribution of eumelanin and phaeomelanin across the body of the dog and along the individual hair shafts, as dictated by a four-allele dominance hierarchy: *a^y^* (fawn) > *a^w^* (wolf sable) > *a^t^* (tan points) > *a* (recessive black) (3–5). However, regardless of the accompanying *ASIP* genotype, the *Melanocortin 1 Receptor* (*MC1R*) gene determines the ability of a melanocyte to produce eumelanin at all, presenting its own dominance hierarchy of four alleles: *E^M^* (melanistic mask) > *E^G^* (grizzle/domino) > *E* (wild type) > *e* (recessive red) (6–8). While the *E^M^*, *E^G^*, and *E* alleles all allow for the production of both eumelanin and phaeomelanin, dependent on the pattern dictated by *ASIP*, homozygous inheritance of *e* prevents eumelanin production, resulting in a completely phaeomelanin color (6–8). Further, the dominant derived *K^B^* allele of *Canine Beta-Defensin 103* (*CBD103*) prevents the production of patterning by *ASIP*, resulting in a solid eumelanin phenotype when combined with a dominant *MC1R* genotype, or a solid phaeomelanin phenotype when combined with a *MC1R* genotype of *e/e* (9). In this way, inheritance of homozygous *e* at *MC1R* or dominant *K^B^* at *CBD103* are epistatic to any of the *ASIP* phenotypes (8,9). In addition to the dominant *K^B^* and recessive *k^y^* alleles of *CBD103*, an intermediate allele, *k^br^*, produces the brindle coat color pattern with alternating stripes of eumelanin and phaeomelanin (10). However, due to the complex structural nature of the *k^br^* allele, which has never been reliably defined and remains unpublished, it was not possible to distinguish between it and *K^B^* for the purposes of this paper.

The base phaeomelanin and eumelanin pigments can be altered by a number of modifier genes, of which *Tyrosinase Related Protein 1* (*TYRP1*), *Proteosome Subunit Beta 7* (*PSMB7*), *Microophthalmia-associated Transcription Factor* (*MITF*), and *RALY heterogeneous nuclear ribonucleoprotein* (*RALY*) are explored further in the scope of this paper. *TYRP1* alters all eumelanin in hair and skin evenly from black to brown, and presents as a compound heterozygote with at least four alternate recessive alleles: *b^s^*, *b^c^*, *b^d^*, and a variant that has thus far only been described in the Australian Shepherd (11–13). *PSMB7* expression produces the harlequin pattern of white, dilute, and deeply pigmented patches only when inherited along with the merle phenotype of *Premelanosome 17* (*PMEL17*, not tested in the present study) (14–16). The *harlequin* variant is homozygous lethal, so the pigmentation phenotype is only expressed when inherited heterozygously (14). *MITF* expression produces white spotting on top of a regularly pigmented background and, depending on the breed background involved, is inherited as a co-dominant or recessive phenotype (17,18). Expression of *RALY*, in combination with other yet-unknown genetic modifiers, alters a tan point background pattern, produced by the *a^t^* allele of *ASIP*, such that the tan points, normally restricted to the paws, muzzle, eyebrows, and chest, extend up the extremities, forming a eumelanin “saddle-shaped” pattern on the dog’s dorsal surface (19).

Expanding beyond pigmentation patterns, other phenotypic traits that are controlled by a relatively small number of genes and can be easily visualized include hair length and texture, tail length, muzzle length, and ear shape. Three genes, *Fibroblast Growth Factor 5* (*FGF5*), *R-spondin 2* (*RSPO2*, not tested in the present study), and *Keratin 71* (*KRT71*), define much of the coat type variation between dog breeds (20–24). Specifically, expression of recessive *FGF5* variants produce a long coat, a dominant *RSPO2* variant produces longer hair specifically on the muzzle and eyebrows, and at least two variants in *KRT71*, one of which was tested in this study, produce a curly coat (20–24). The tailless trait, whereby a dog is born with a truncated or absent tail, is caused in some breeds by a variant in the *T brachyury transcription factor* (*T*) gene (25,26). Like harlequin, taillessness is homozygous lethal, so will be expressed viably only in heterozygotes (25,26). Ear set and muzzle length are traits that are very likely impacted by numerous genetic variants, however a marker on canine chromosome 10 is known to segregate in some cases for erect versus drop ears (27,28), and the *Bone Morphogenic Protein 3* (*BMP3*) gene is associated with foreshortening of the face (29). A variant in *SMOC2* has also been shown to contribute to muzzle length, though it is not included in the present study (30).

Epistatic effects are prevalent within coat color and morphological variation in the dog. We have previously mentioned epistasis between a homozygous non-functional genotype at *MC1R* or a dominant variation at *CBD103* with the ability to express *ASIP*-driven phenotypes. However, canine coat color also presents scenarios whereby a specific allelic background is required for expression of a modifier. This is exemplified in the requirement of the merle phenotype of *PMEL17* in order to express the harlequin pattern of *PSMB7*, the necessity for an *a^t^* tan point base pattern at *ASIP* to display the *MC1R* grizzle or *RALY* saddle tan phenotypes, and a moderate to long hair length to produce a *KRT71* curly coated phenotype. On an incompatible background, presence of some gene variants may remain unexpressed for generations or, as many breeds are fixed for only a small number of phenotype options, may remain completely unobserved. As national and international breed organizations are charged with defining their breed’s characteristics, which are then regulated by registering bodies such as the American Kennel Club (AKC), United Kennel Club (UKC), The Kennel Club (KC) in the United Kingdom, or Fédération Cynologique Internationale (FCI), failure of breed standards to account for rare - though naturally occurring - variation can lead to frustration or confusion when unexpected traits are expressed. That same existence of rare variants within breeds – which, due to epistasis and genetic background, may never express the associated phenotypes - can provide intriguing information regarding the development of, or relationships between, breeds throughout history.

We utilized custom genotyping array data from Mars Wisdom Health for 11,790 canids, representing 212 pure breeds and 4 wild canine populations (S1 Table), genotyped for seven coat color and five physical characteristic genes (Table 1). For each of these genes, we determined the frequency of each allele within each breed. Note that for one gene (*TYRP1*) not all alleles could be tested, these are indicated in Table 1. Utilizing these frequencies, we then evaluated: 1) the breed-type distribution of morphologic variants; 2) implications of founder effects and/or selection preferences between geographically or behaviorally independent populations of the same breed; 3) the breed-specific carrier status of variants disallowed within breed standard descriptions; and 4) the ancestral connections between breeds that share rare trait-causing variants.

**Table 1:**
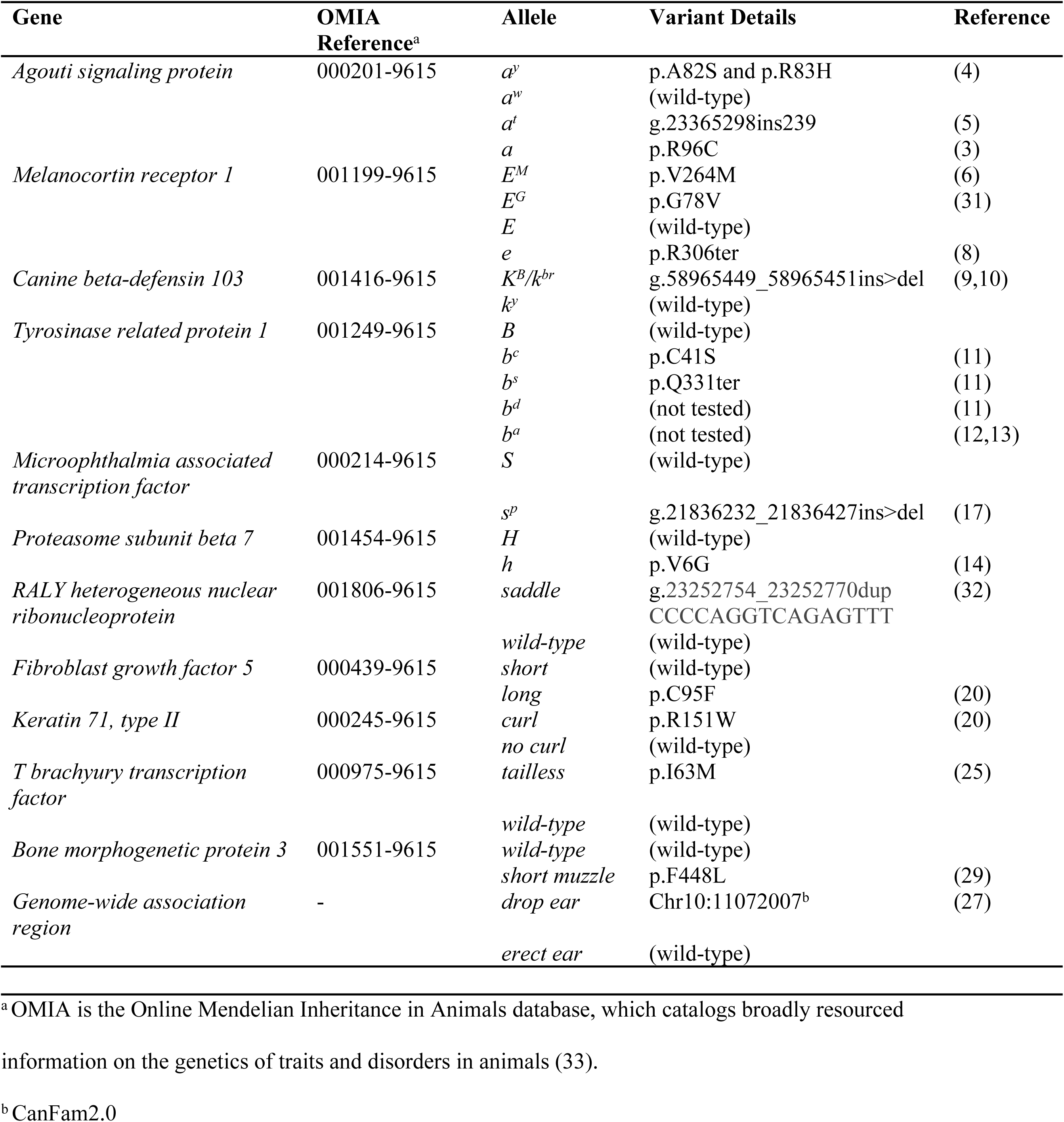
Genes and markers genotyped for breed analysis.

## Results

### Unexpectedly broad distribution of trait-causing alleles across breeds

Marker genotypes were combined to interpret the actual biallelic genotypes for each of the queried genes for every dog in the dataset. Individuals of the same breed were combined to calculate breed allele frequencies and are reported in S2 Table. Breeds were assigned to phylogenetic clades as previously reported (34,35)(S1 Table). Breeds not included in the earlier phylogenetic studies were assigned to defined clades based on known breed history, phenotypic commonalities, and geographic region of origin (indicated by parentheses in S1 Table).

Over all combined breeds, the ancestral allele (Table 1) predominates at all genes, except for *ASIP* and *MC1R*, where derived alleles account for 82% and 57% of alleles, respectively. The derived alleles at lowest frequency across all 11,790 canids are *E^G^* (grizzle/domino) of *MC1R* and *tailless* of *T*, each representing ∼1% of all alleles detected within their respective genes. *E^G^*, which produces the grizzle or domino pattern when combined with *a^t^* tan points at *ASIP*, is present in 28 breeds, with highest frequencies recorded in Borzois (56%), Polish Greyhounds (43%), and Salukis (31%). Cladistic representation of the *E^G^* allele is heavily weighted to those breeds in the Mediterranean and UK Rural clades (Figure 1), with 60% and 31% of all *E^G^* alleles in our dataset arising from those clades, respectively. The presence of the *E^G^* allele has previously been reported in only the Saluki and Afghan Hound (31), therefore, our analyses document the existence of the allele in an additional 26 dog breeds (S2 Table).

**Fig 1:**
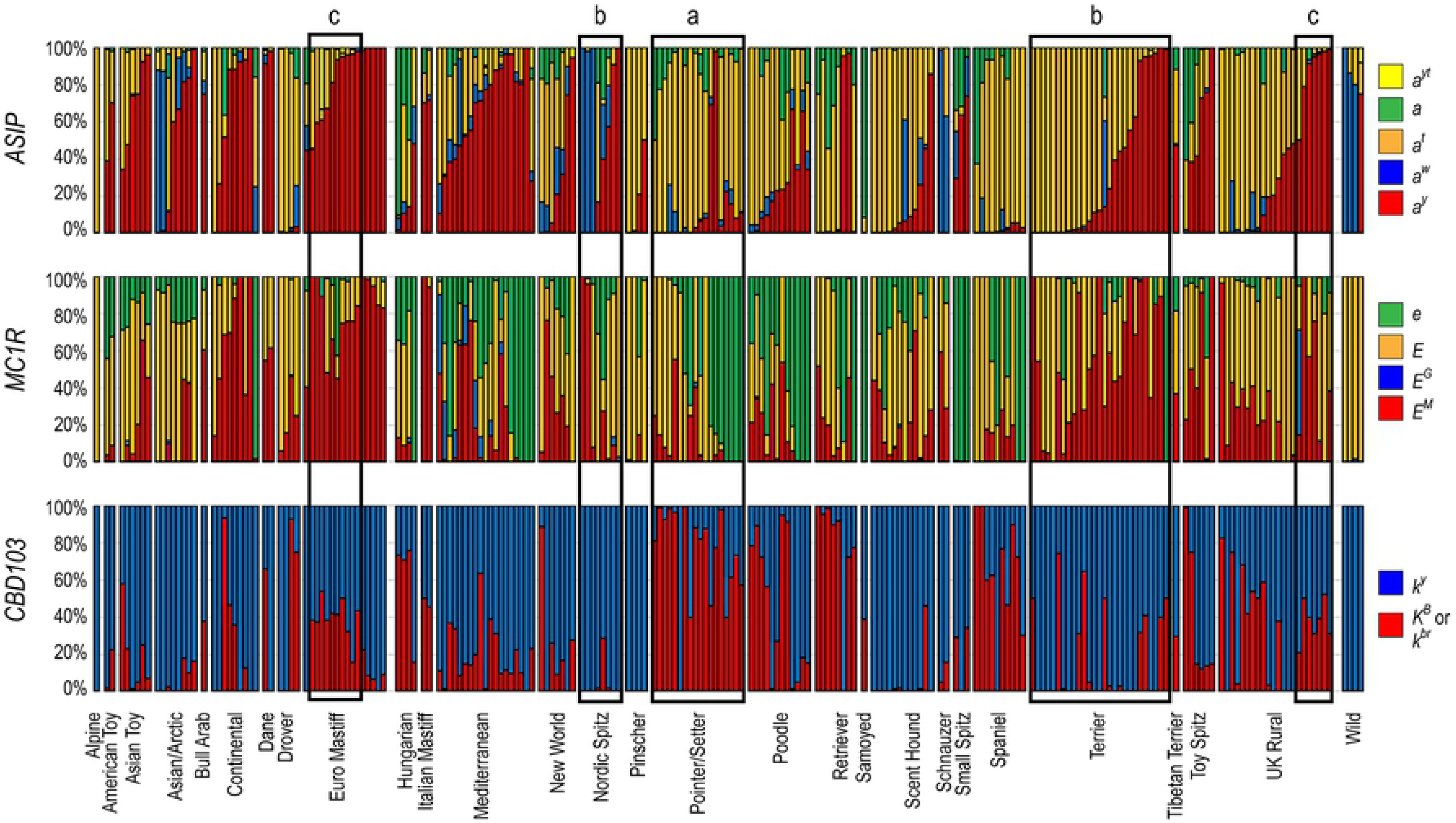
Allele frequencies for *ASIP*, *MC1R*, and *CBD103* by phylogenetic breed relationship. Breeds are grouped by phylogenetic clade, as described, and sorted within clade by frequency of *e* and then *a^y^* to demonstrate patterns of phenotype expression across interacting genes. Thick black boxes highlight examples of color preference influencing interacting genes: a) Pointer/Setter breeds are commonly seen in solid colors, caused by the *K^B^* allele of *CBD103* or the *e*/*e* genotype of *MC1R*. Since the solid-color genotypes are epistatic to the dominant alleles of *MC1R* and all alleles of *ASIP*, variation at those genes does not follow a trend for color preference. b) Related breeds with high frequency of the *k^y^* allele of *CBD103* have a more structured pattern of *ASIP* allele preference. c) Breeds with a preference for the brindle pattern show heterogeneity for *K^B^*/*k^br^* (reflective of a *k^br^* phenotype) and high frequency of *a^y^*, required for the expression of brindle across the whole body.

Assignment of biallelic genotypes for the *ASIP* gene based on genotype tests for *a^y^*, *a^t^*, and *a*, and a genotype of *a^w^* by exclusion of the other three alleles, revealed the presence of unusual triple-allele combinations in a few dogs. In these cases, dogs would appear to genotype as *a^y^*/*a^t^*, plus an additional third allele. This genotyping outcome reflects a scenario where the *a^y^* point mutations and the *a^t^* SINE insertion occur on the same chromosome. The resultant combinatorial allele, termed *a^yt^*, was identified in 14 domestic breeds across 10 clades, and the Dingo (S2 Table). Phenotype descriptions of the evaluated dogs were not available, so the effect of the *a^yt^* allele on phenotype is presently unknown. The breed with the highest frequency of *a^yt^* is the Dogo Argentino (33%), a predominantly white mastiff breed.

The *RALY* duplication required for the *saddle* modification of the tan point phenotype was detected in a greater than expected number of breeds (n = 203) (S2 Table). Based on breed standard descriptions, 17 of the breeds observed to have the *RALY* duplication should be at or near fixation for the saddle tan phenotype. Indeed, these breeds have a *saddle* allele frequency of between 76-100% (average = 95%) and an *a^t^* tan point allele frequency between 38-100% (average = 89%), supporting the association of the saddle tan phenotype with the combined genotypes of *ASIP a^t^* and the *RALY* duplication. Both the tan point and saddle tan phenotypes are allowed and observed in a further 21 breeds, with observed *saddle* frequencies between 24-98%, and *a^t^* frequencies of 15-94%. Fourteen breeds show appreciable levels of *a^t^* frequencies (44-100%) and *saddle* frequencies (8-100%), without regular expression of the saddle tan phenotype within the breed.

The *tailless* allele was detected in 48 breeds, with greatest frequencies in the Tenterfield Terrier (30%), Swedish Vallhund (18%), Spanish Water Dog (14%) and Australian Shepherd (13%). The remaining 44 breeds carry the *tailless* allele at <10%. The breeds that are suggested to carry *tailless* are widely distributed across 14 phylogenetic clades, but are more heavily present in the UK Rural (9 breeds), Terrier (8 breeds), Poodle (6 breeds), European Mastiff (4 breeds), and Pointer/Setter (4 breeds) clades (S1 Figure). Previous reports have documented the *tailless* allele in 18 breeds (25,26), ten of which overlap with our dataset and likewise display the *tailless* allele (the remaining eight are rare, regional breeds and not included in the present dataset). The current dataset also includes the suggested presence of the *tailless* allele in an additional 38 breeds, in which the allele has not been previously reported.

The recessive *a* allele of *ASIP* has previously been reported in six breeds (German Shepherd Dog, Belgian Shepherd, Schipperke, Australian Shepherd, Shetland Sheepdog, and Eurasier) (3–5). We identified the allele in 89 breeds, documenting for the first time the presence of the allele in an additional 83 breeds (S2 Table). The allele was observed in 23 clades. The clades with the highest number of *a*-possessing breeds are the Pointer/Setter (13 breeds), Poodle (8 breeds), Spaniel (7 breeds), and UK Rural (7 breeds) clades.

Twenty-six breeds are fixed for a single allele at *ASIP* as follows: *a^y^* 9 breeds, *a^w^* 2 breeds, and *a^t^* 15 breeds. Thirty-nine breeds are fixed for a single allele at *MC1R* as follows: *E^M^* 13 breeds, *E* 9 breeds, *e* 17 breeds (S2 Table). No single breed is fixed for one allele at every gene evaluated. Four populations of wild canines were included in the analyses, although they are not counted as “breeds” in the analyses presented here. The wild canine populations had high levels of wild-type fixation across all genes evaluated (S2 Table). Eastern coyotes (n = 29) show a 2% frequency of the *MC1R E^G^* allele and a 2% frequency of the *TYRP1 b^s^* allele. The Western coyotes (n = 19) have a 21% frequency of the *FGF5 long* allele. The Dingo samples (n = 12) show an 8% frequency of the *TYRP1 b^s^* allele. The marker for ear shape, and the causal variants in *RALY*, *MITF*, and *ASIP* are moderately variable across all four wild populations.

### Cladistic patterns of allele distribution

Allele frequencies of *ASIP*, *MC1R*, and *CBD103* are graphically represented in Figure 1, relative to the phylogenetic groupings of the breeds. Association of allele frequency with cladistic assignment highlights multiple key tendencies in regards to color preference within breed-type groups. The cladistic distribution of *CBD103* alleles reflects a tendency toward fixation of a single allele within related breeds. The Hungarian, Pointer/Setter, Poodle, and Retriever clades contain breeds nearing fixation for the *K^B^/k^br^* variant, presumably primarily *K^B^* due to the predominantly solid-colored phenotypes of breeds in these clades. Conversely, the Asian/Arctic, Mediterranean, New World, Scent Hound, and Terrier clades consist of breeds nearing fixation for *k^y^*. Since the *K^B^* allele produces a solid colored animal and is epistatic to alleles of *ASIP* and all *MC1R* alleles except for *e*, variation of *MC1R* and *ASIP* alleles is relatively uncontrolled in breeds with a high frequency of *K^B^*. Comparatively, clades with high frequency of the *k^y^* allele show greater degrees of preference for the patterning alleles of *ASIP*, specifically *a^y^*, *a^w^*, and *a^t^*. Clades with strong heterogeneity of *CBD103*, represented as multiple breeds within a clade with *K^B^* frequency of 40-50% (eg. Euro Mastiff, portions of UK Rural), are also those that consist of breeds presenting the brindle coloration. This is an accurate reflection of the present inability to distinguish between the brindle *k^br^* and dominant black *K^B^* alleles due to the complex genetic structure of the variants. Comparatively, clades with multiple breeds that show a 40-50% frequency of *K^B^* likewise have a high frequency of *a^y^*, the *ASIP* background that is necessary to produce brindling across the entire body of the dog.

The wild-type *a^w^* allele of *ASIP*, while predominant in three of the four wild canine populations, is at greatest frequency in the spitz clades (Asian/Arctic, Nordic Spitz, Schnauzer). For *MC1R*, the *e* allele was identified in 24 of the 28 clades represented in this data, but it is seen at highest frequencies in hunting breeds (Pointer/Setter, Poodle, Retriever, Spaniel clades), continental sighthounds and flock guardians (Hungarian, Mediterranean clades), and white spitz breeds (Samoyed, Small Spitz clades). The melanistic mask allele, *E^M^*, was likewise widespread, observed here in 164 breeds across 24 clades. Finally, the *E^G^* allele is most prevalent in the Mediterranean clade, represented only sporadically in 11 other clades.

Allele frequencies for the nine remaining genes tested, sorted relative to phylogenetic breed relationships, are presented in S1 Figure. With the exception of the harlequin mutation, detected only in Great Dane and Yorkshire Terrier, all remaining gene variants were present broadly across all clades. There is no apparent excess of specific *TYRP1* brown alleles in any given clade or breed, despite, or perhaps due to, the identical phenotype produced by both alleles. Breeds fixed for the *MITF* SINE insertion, associated with piebald white spotting, are most abundant in the Pointer/Setter and Terrier clades, but are still widely distributed. Similarly, the *saddle* variant of *RALY* is seen in numerous clades, but is most abundant in the Euro Mastiff, Mediterranean, Scent Hound, and Terrier clades. Only eight breeds (AIRT, BEDT, IWSP, KOMO, LAKE, PULI, WELT, WFOX), in the Hungarian, Retriever, and Terrier clades are entirely fixed for the hair *curl* variant of *KRT71*. The *long* haired variant of *FGF5* is at the highest levels in the American Toy, Asian Toy, Continental, Hungarian, Poodle, Retriever, Small Spitz, and Spaniel clades. Conversely, the highest representation of the *short* haired wild-type genotype is in the Euro Mastiff clade. Toy breeds, Mastiffs, and Terriers represent the highest frequencies of the *BMP3 short muzzle* genotype. Finally, the only clades with breeds that are fixed for the *drop* ear marker are the Pointer/Setter, Retriever, Scent Hound, and Spaniel clades.

### Allele frequency is influenced by within-breed selection and geographic separation

Thirty breeds were divided into two or more distinct populations based on geographic region (27 breeds), body size (Dachshund and Poodle), coat type (Dachshund), and lineage or application, such as those selected for working applications (i.e. field, racing) or conformational traits (i.e. show) (6 breeds). All such populations were genetically distinguishable via principle components analysis (PCA). For the present study, allele distributions of the 12 tested genes between same-breed populations were evaluated for significance using either Pearson *χ*^2^ or Fisher’s Exact test (the latter was used when any allele grouping presented at <5 individuals). Distributions were deemed statistically significantly different at p ≤ 0.00167, after correction for multiple testing. Nine breeds (BASS, BULM, DALM, KEES, MANT, POM, SCWT, WEIM, WELT) showed no significant difference between populations in allele distribution at any evaluated gene. The remaining 21 breeds had significant allele distribution differences between populations for at least one gene (Figure 2, S2 Figure). The gene with the greatest number of breeds displaying differences in allele distribution by population is *MC1R*, which was statistically different in 11 breeds (Figure 2). The *T* gene, responsible for the *tailless* phenotype, is not statistically different between populations of any breed (S2 Figure).

**Figure 2:**
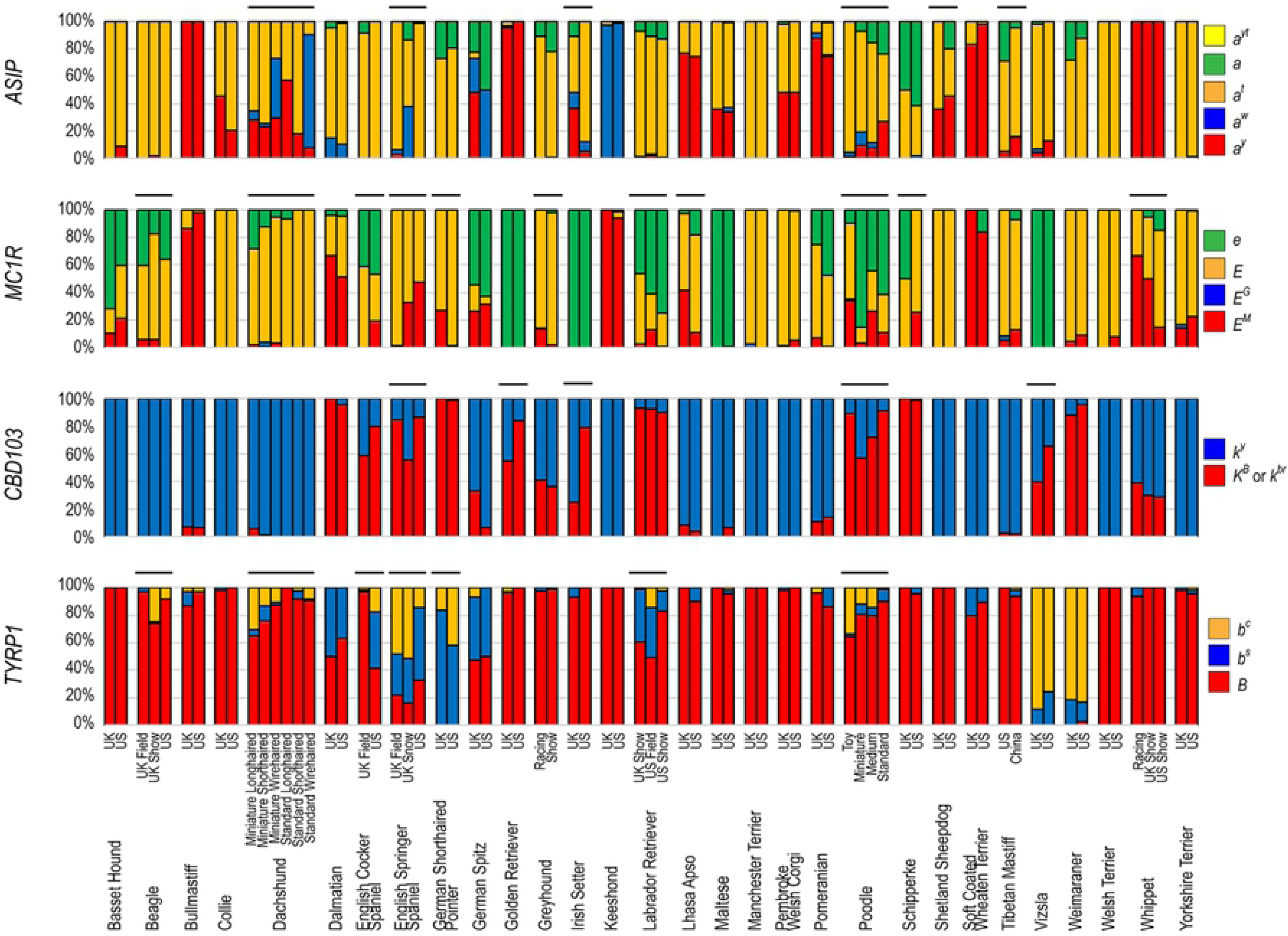
Allele frequencies for *MC1R*, *ASIP*, *TYRP1*, and *CBD103* across same-breed populations. Horizontal black bars indicate within-breed allele distributions that are significantly different (p ≤ 0.00167).

The effect of same-breed population divergence is readily observable in the system of coat color gene interactions and epistasis. Five breeds (ESSP, GOLD, ISET, POOD, VIZS) have significantly different *CBD103* allele distributions between same-breed sample populations (Figure 2). Of these five breeds, three are uniformly fixed for an *MC1R* genotype of *e*/*e* (GOLD, ISET, VIZS), effectively preventing the expression of the phenotype variability that the different alleles of *CBD103* would otherwise produce. Conversely, regional phenotype preference can be seen in breeds such as the Labrador Retriever (Figure 2). The breed shows significantly different allele distributions by population for *MC1R* and *TYRP1*, indicating that US conformation show lines prefer yellow (*MC1R e*/*e*) dogs with black noses (*TYRP1 B*/_), UK conformation show lines prefer black (*MC1R E*/_, *TYRP1 B*/_) dogs, and US field lines consist mainly of brown (*MC1R E*/_, *TYRP1 b*/*b*) or yellow dogs (*MC1R e*/*e*) (p ≤ 0.003).

### The natural probability of disallowed traits in pure breeds

The approved phenotypes for each breed were determined based on the written breed standards of the AKC, FCI, UKC, and KC (Britain). Of the 212 breeds evaluated, 143 were observed to carry at least one allele that would result in an unfavorable phenotype, termed “fault”, relative to at least one of the four queried breed registries (S3 Table). The breeds with the highest number of fault-causing alleles are the Treeing Walker Coonhound and Great Dane with 6 fault alleles present in each, and the Yorkshire Terrier and Rottweiler with 5 fault alleles present in each. Seventy-eight percent of the total number of fault allele occurrences produce phenotypes disallowed by all queried registries that recognize the breed. For 8.9% of fault allele observations, one or more of the queried breed registries allow the phenotype produced without bias, with the remaining registries disallowing it entirely or imparting a preference bias. In 15.1% of the fault allele observations, at least one of the four registries describes the allowed phenotypes indistinctly. This consists of situations where the trait is preferred to a lesser extent than alternatives, is not described at all, or is worded in such a way as to lead to ambiguous interpretation. The most frequently observed fault-causing alleles are the recessive brown alleles of *TYRP1*, representing 29.8% of all fault allele instances. Notably, 79.6% of all fault alleles are recessively inherited.

The probability of producing the disallowed phenotype associated with each fault allele was calculated assuming a random breeding same-breed population, and taking into consideration the inheritance patterns and gene interactions required for the phenotype expression. The fault-producing probabilities range from 4.9e^−7^ (any non-solid color in the Black Russian Terrier) to 0.25 (red in the UK population of Schipperkes). Overall, the fault alleles were detected at low frequencies and, due to complex inheritance hierarchies and epistasis, have a <0.01 probability (<1% chance) of producing the fault phenotype in 58.9% of instances. Only 4.2% of fault alleles have a >0.10 probability (>10% chance) of producing the fault phenotype.

### Ancestral routes of allele transmission

Previous research has demonstrated the ability of identity-by-decent (IBD) haplotype sharing to reveal shared ancestry events between modern dog breeds, accurate to approximately 150 years ago (34). We used these previously identified significant breed relationships to successfully connect breeds genotyped to possess the rare phenotype alleles: *T tailless* (Figure 3) and *ASIP a* (Figure 4). Forty-eight breeds were identified as carrying the *tailless* allele of *T*, in many cases at very low frequencies, in the present study. Thirty-eight of these breeds were represented in previous research (34,35), allowing identification of IBD haplotype sharing relationships. By using these pre-determined breed relationships, each of the 38 breeds (Figure 3, represented by red or green text) can be connected into a single relationship matrix, providing a potential route of transmission of the *tailless* allele. Only nine non-carrier breeds (Figure 3, represented by grey text) were required to serve as potential transmitters of the *tailless* allele, reflecting populations that have successfully eliminated the allele from the breed, or have decreased the frequency enough so as to not be detected in our sampling. Likewise, 89 breeds were identified as carrying the *ASIP a* allele in the present study, 68 of which were also analyzed for IBD haplotype sharing previously (34,35). Of these 68, all but three breeds (Tibetan Mastiff, Kuvasz, and Anatolian Shepherd) could be connected via significant IBD haplotype sharing events (Figure 4).

**Figure 3:**
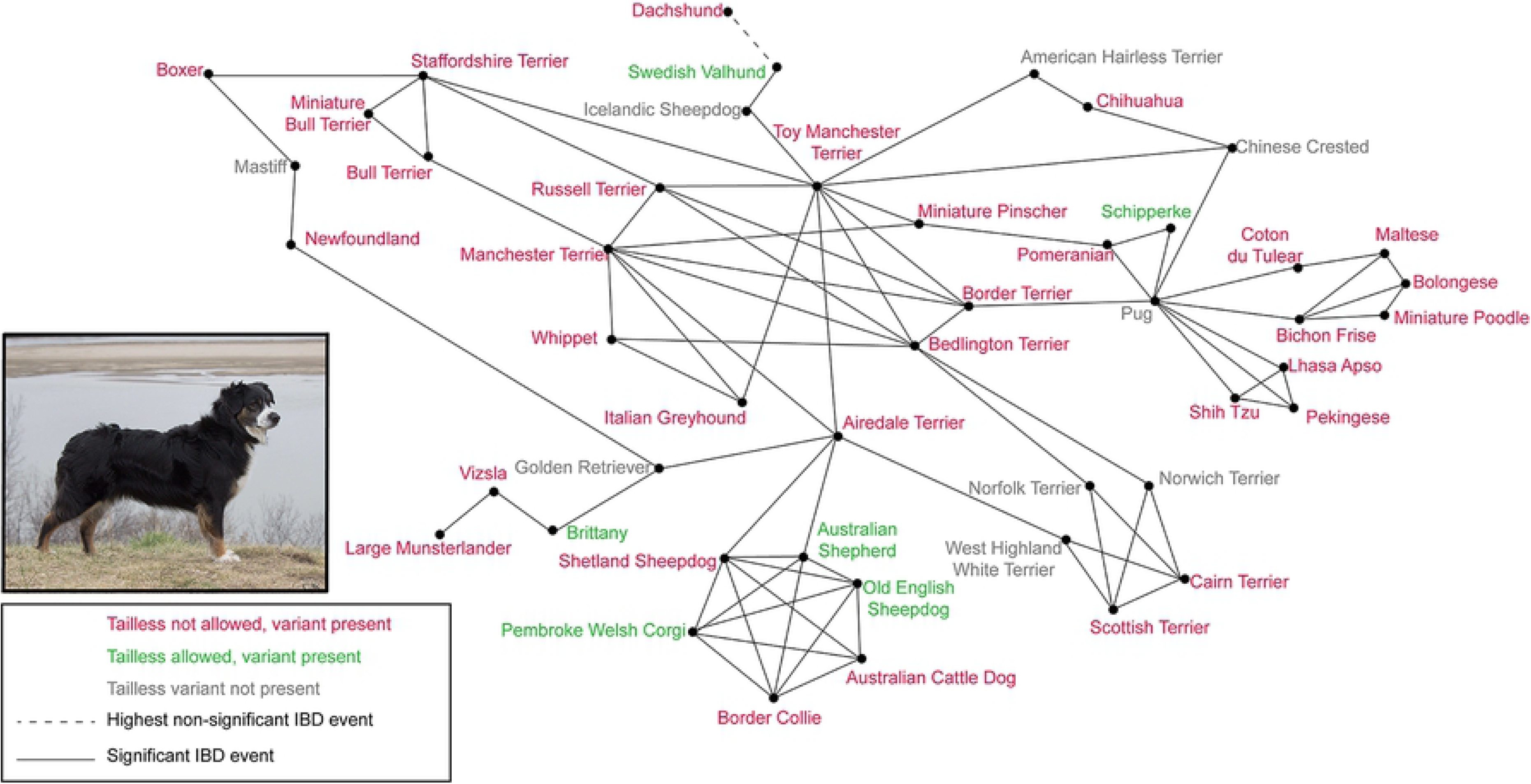
Identity-by-decent (IBD) haplotype sharing breed relationships connect breeds carrying the *T* allele for *tailless*. Solid black lines represent instances of significant haplotype sharing levels between breeds. The color of the breed names reflects the proposed carrier status of the tailless variant in the sampled members of those breeds, indicating not present (grey), present and permitted within the breed standard (green), and present and not permitted within the breed standard (red). The Dachshund breed shows no significant haplotype sharing with any other breed, however, its highest non-significant haplotype sharing value is with the Swedish Valhund (dashed line). Inset, the Australian Shepherd breed permits natural taillessness.

**Figure 4:**
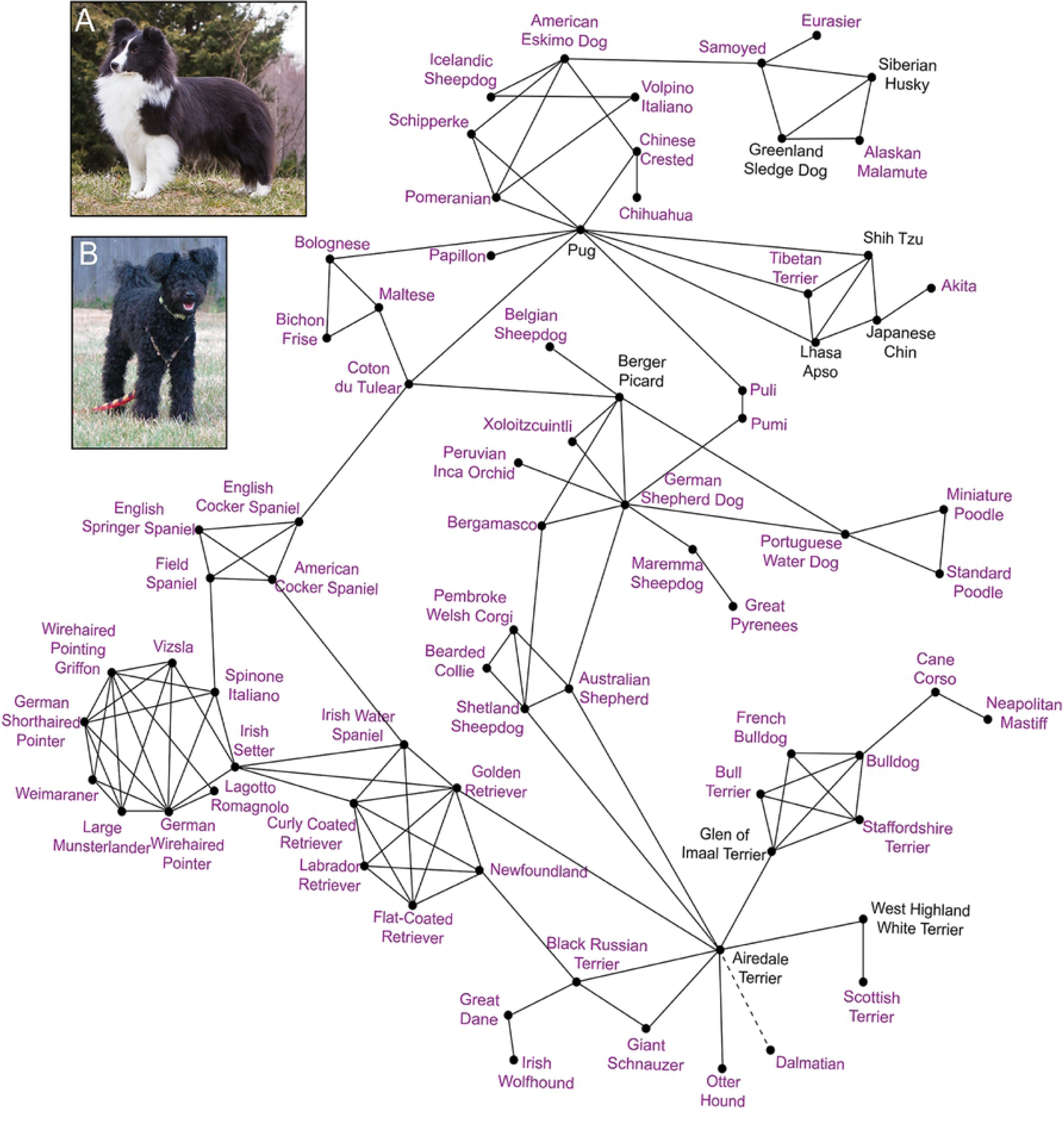
Identity-by-decent (IBD) haplotype sharing breed relationships connect breeds carrying the *a* allele of *ASIP*. Solid black lines indicate instances of significant haplotype sharing between breeds. Breed names in purple indicate observed frequency of the *a* allele within the breed in the present study. Breed names in black indicate that the *a* allele was not detected within the sampled individuals of that breed. The dashed line connecting the Dalmatian to the Airedale Terrier indicates that no significant haplotype sharing was detected with the Dalmatian, however the highest non-significant level of haplotype sharing was measured with the Airedale Terrier. Inset images show examples of dogs with the recessive black phenotype, A) Shetland Sheepdog, B) Pumi.

## Discussion

Most publications to date that assess allele frequency within dog breeds have been conducted on a relatively small scale, limited to select sets of breeds and/or variants of known relevance to specific breeds (36–42). A recent publication focused on a very large set of purebred and mixed-breed dogs and a panel of 152 disease markers (43), but did not evaluate morphological or pigmentation gene variants. Here, we have leveraged an expansive number of DNA samples, both in terms of dog numbers and breed representation for which 12 genes impacting coat color and morphological variation have been genotyped on a commercial genotyping platform (Table 1). In doing so, we have revealed patterns of allele frequency and distribution that inform our understanding of breed development and relationships, regional phenotypic preferences, and the impact of canine registering bodies on the prevalence of rare alleles.

### Unexpected breed distribution of low frequency alleles

The majority of the alleles evaluated in the present study are broadly recognized across many modern dog breeds. However, a small number of the variants have only previously been recognized in select breeds. For example, the *E^G^* allele of *MC1R* was initially detected in Salukis and Afghan Hounds, due to the popularity of a unique color pattern in those breeds, termed grizzle or domino, respectively (7). We have here described 26 additional breeds in which the *E^G^* allele has not previously been reported (S2 Table). Related sighthound breeds can have patterns similar to grizzle/domino, such as “sable” in the Borzoi, or “wolf” in Polish Greyhounds, and unsurprisingly they were found to carry the *E^G^* allele. Seven additional breeds (ANAT, CAAN, CASD, CAUC, KKLG, POLG, TAIG) carrying *E^G^* are phylogenetically related to the Saluki and Afghan Hound, suggesting shared traits due to common ancestry, but *E^G^* was also found in the Asian/Arctic, Hungarian, Nordic Spitz, Pointer/Setter, Poodle, and Scent Hound clades. The *E^G^* allele requires an *ASIP* tan point phenotype in order to express as an observable pattern (7), the lack of which would allow the allele to persist within a breed undetected. This may be the case in breeds such as the Anatolian Shepherd, Black Russian Terrier, Maltese, Norwegian Buhund, Puli, and Toy Poodle, where the tan point phenotype is rare or absent.

While not necessarily considered a low-frequency allele, the *E^M^* “melanistic mask” allele of *MC1R* warrants discussion. Despite anecdotal observation in numerous breeds, previous publications have identified the *E^M^* allele in only 11 breeds (6,31). We have identified the allele as present in 164 breeds, 153 not previously documented. In at least 38 of these breeds, the mask phenotype is not observable, through epistatic interactions with non-compatible genotypes such as solid eumelanin (eg. DALM, GSHP, LMUN, WEIM, PTWD, NEWF, SKIP, OES) or solid white (DOGO, BBLS, COTO) coat colors.

The *tailless* allele of the *T* gene has previously been investigated in breeds known to have the natural tailless phenotype, confirming the cause of the trait in 18 breeds (25,26). The frequency of the variant has not, however, been evaluated in breeds not necessarily expected to harbor the trait. We report the occurrence of the *tailless* allele in 38 new dog breeds (S2 Table, Figure 3). The tailless phenotype is variably expressed, ranging from complete anury to a truncated or kinked tail (25,26). While absence of a tail in a breed expected to have a full length tail would be immediately noticeable, a mildly shortened or bent tail may be dismissed as inconsequential, thus allowing a low frequency of the allele to persist within some breeds. Likewise, surgical tail docking is still a common practice among certain breeds in some countries, rendering the natural length of the tail unknown beyond a neonatal age. Among the 38 newly reported carrier breeds of *tailless*: two allow a natural tailless phenotype (MCNB, OES); seven follow tail docking procedures in the US (AIRT, BOX, MPIN, MPOO, TENT, VIZS, WELT); eight have breed standards that describe a shorter than average ideal tail length (BORT, BULT, CAIR, MANT, MBLT, RWST, SCOT, TMNT); and nine have naturally curled tails (BICH, BOLO, COTO, GSPZ, LHAS, MALT, PEKE, POM, SHIH), which would mask small variations in length or structure. Indeed, anecdotal reports of abnormally short and/or kinked tails have been reported in at least eight of the newly reported *tailless* carrier breeds (BEDT, BULT, DACH, LHAS, SSHP, STAF, SCOT, VIZS) (44–48)(personal communications with breeders). It is important to note that many of the breeds in which we have now detected the *tailless* allele show the allele as present at very low frequencies. This allows for the possibility that some of these calls may be false positive errors that have escaped our quality control efforts.

The identification of an allele with the combined *a^y^* and *a^t^* variants in the *ASIP* gene poses a number of intriguing implications. The variant associated with the *a^t^* allele is a SINE insertion located on CFA chr24:23,365,298-23,365,537, in the upstream regulatory region of *ASIP,* and has been attributed to dorsoventral and banded hair patterning (5). The *a^y^* variant consists of two adjacent amino acid substitutions in exon four of the *ASIP* gene (p.A82S R83H), located at CFA chr24:23,393,510-23,393,515 (4). At approximately 28 kb apart, a canine recombination rate of 1.2 cM/Mb (49) would predict a 0.042% rate of crossover between the two variants. We observed the *a^y^* and *a^t^* variants on the same chromosome, briefly termed the *a^yt^* allele, a total of 48 times within the 11,790 genotyped dogs, and within 14 seemingly unrelated breeds, and the Dingo. Three of these breeds, the Maremma Sheepdog, Great Pyrenees, and Dogo Argentino, are predominantly white in color, masking any pigmentation variation that may be present due to the *a^yt^* allele. The phenotypes of individual genotyped dogs were not available, but relying on the accepted colorations with each breed, the remaining 11 breeds with *a^yt^* all allow for the standard *a^y^* fawn phenotype. Frequencies of *a^yt^* varied within the fawn breeds from 1% to 8%, suggesting that any phenotypic difference caused by the allele combination would not be immediately recognized as outside the acceptable *a^y^* fawn coloration. Indeed, with the added *a^t^* SINE insertion, preventing the production of banded hair patterning, *a^yt^* may result in a fully phaeomelanin color, lacking the eumelanin tips on the hairs that are commonly present in *a^y^* fawn dogs. The current method of determining the biallelic genotype of *ASIP* relies on assaying the *a^y^*, *a^t^*, and *a* variants, and assigning the presence of *a^w^* in the absence of the other detectable markers. In this way, a genotype of *a^yt^*/*a^w^* and *a^y^*/*a^t^* would appear to be the same. Seven breeds (ANAT, DANE, GPYR, LAGO, MARM, TIBM, TIBS) in which the *a^yt^* allele was detected also showed observable levels of the *a^w^* allele, presenting situations where the true frequencies of *a^w^* may be higher than reported here. An additional 63 breeds have the *a^y^*, *a^w^*, and *a^t^* alleles, reflecting the opportunity for the *a^yt^*/*a^w^* genotype to be incorrectly recorded as *a^y^*/*a^t^*, which would also result in underestimate of *a^w^* frequency. There is, of course, the possibility that either the *a^y^* or *a^t^* variants previously described are not causal variants, but rather very accurate markers. However, both variants have been successfully used to predict coat color genotypes in multiple studies without raising concern (7,32,50,51). Further pursuit of the phenotypic impact of the *a^yt^* allele, along with determining its inheritance pattern, is ongoing.

The *RALY* duplication associated with the saddle tan modification of the tan point phenotype was detected in 203 breeds (S2 Table). Thirty-eight of these breeds are either fixed or variable for the saddle tan phenotype, while 119 of these breeds have epistatic variation at additional coat color genes that prevent the expression of *saddle tan*. However, both the saddle tan and tan point mutations were detected in 14 breeds for which only a tan point phenotype is permitted or expected. For instance, Bernese Mountain Dogs have a fixed phenotype of black, tan points, and white spotting; despite the 100% *a^t^* allele frequency and the addition of an 18% *saddle tan* allele frequency, no readily identifiable dogs exist with the saddle tan phenotype. The remaining 32 breeds have breed standards that would permit for the occurrence of a saddle phenotype, but the pattern is rarely observed due to low frequency of the required tan point genotype. These instances reiterate the findings of the initial canine *RALY* research, suggesting at least one additional modifier gene that is required for the production of the saddle tan phenotype (32).

The *Harlequin* allele was identified, as expected, in the Great Dane population (14), but was also identified – unexpectedly – in the Yorkshire Terrier population, where it has not been previously described. Yorkshire Terriers are not known to possess the variant at *PMEL17* that produces the merle phenotype and is required for the expression of harlequin when combined with the *PSMB7* variant. Therefore, even with the *Harlequin* mutation, it would presently be impossible to produce a harlequin patterned Yorkshire Terrier. Despite this, there is no recent haplotype sharing between the Yorkshire Terrier and Great Dane breeds, which could have resulted in transfer of the allele (34). The detected presence of the allele in our data set could be explained by a *de novo* mutation in the Yorkshire Terrier breed, inaccurate marker selection, or genotype quality in the SNP array. The latter is unlikely due to the accuracy of this test on the positive control population (Great Danes) and the negative control population (all other dog breeds). The *de novo* mutation theory is supported by the fact that *Harlequin* was only identified in the UK Yorkshire Terrier population (2 dogs with the *Harlequin* allele out of 24 UK Yorkshire Terriers) and not at all in the US Yorkshire Terrier population (n = 107 dogs); these two populations interbreed only rarely, suggesting the *de novo* mutation arose privately in the UK. Further investigation is required to draw definitive conclusions.

Given the expectation of ancestral alleles in wild canine species, the level of derived alleles detected in the wild canine populations was somewhat surprising. While 99% of *MC1R* alleles in the Grey wolf, Eastern coyote, Western coyote, and Dingo were found to be wild-type *E*, only 72% of *ASIP* alleles from wild canines were wild-type *a^w^*, with only the Western coyote fixed for *a^w^*. The genotyped Dingos were predominantly (75%) *a^y^* fawn, consistent with their usual phenotype. Somewhat unexpectedly, however, the *a^t^* tan point allele of *ASIP* was detected at levels of 20% in Eastern coyotes and 14% in Grey wolves. Likewise, derived coat color alleles at *TYRP1* and *MITF*, and the morphologic variants associated with long hair, and drop ears, were detected at within-species levels between 2-46% (*b^s^* in Eastern coyotes and drop ear in Grey wolves, respectively). The derived alleles at *ASIP*, *MC1R*, *CBD103* have previously been detected at low frequency in various populations of coyotes and wolves, most of which have been postulated to occur due to introgression with domestic dogs (51–54). Such introgression between grey wolves and domestic dogs has been documented to occur at low levels across Eurasia, but detailed studies of domestic dog introgression with wolves and other species (coyotes, dingoes) in other geographic areas are sparse (55,56). Our grey wolf (n = 12) population was largely collected on the East coast of North America, and theoretically has more opportunities for introgression with domestic dogs due to higher human (and pet dog) population density compared to their Western North American counterparts. This is the first report of the tan point (*ASIP*), and long hair (*FGF5*) variants in wild canine populations. A recent paper verified the CFA10 ear shape locus, and demonstrated that, while definitively a contributing locus, it does not perfectly account for ear phenotype (28). Thus, the higher frequency of ear shape and hair length variant alleles in wild populations is perhaps not surprising, as these phenotypes are known to be driven by more than one locus (20,27,54).

### Intended working application influences phenotypic variation

Phylogenetic analysis of dog breeds is conducted in the absence of phenotypic data and provides intriguing insight into the history of breed formation. Reversing that scenario by assessing phenotypic association relative to phylogenetic relationships begins to unravel the individual characteristics that define cladistic groupings. The comparison of allele frequencies of the interacting coat color genes *ASIP*, *MC1R*, and *CBD103* (Figure 1) reveal preferred clade-specific colorations. *ASIP*-driven patterns are widely preferred in the Scent Hound, Spitz, and Terrier clades, specifically tan point, wolf sable, and a split of fawn and tan point, respectively. Conversely, single-pigment patterns, characterized by high frequencies of *CBD103 K^B^* or *MC1R e* alleles, are prevalent in the clades with a history of hunting applications, namely those of the Pointer/Setter, Poodle, Retriever, and Spaniel clades. Similarly, the hunting-related clades also generally display a higher incidence of brown pigment production, though they do not show preference for a specific recessive *TYRP1* allele (S1 Figure).

Historically, white has been selected for in breeds to improve their visibility. Breeds used for hunting served two primary purposes: run ahead of the hunter to locate prey and cause it to fly or run out from hiding, or remain stationary with the hunter until sent to retrieve downed game or lost arrows (57,58). Ability of the hunter to visually locate their dog while it searches for game is said to be aided by the adding of white spotting patterns, as is the case with pointer, setter, and spaniel breeds (58–61). Conversely, when a dog is meant to remain camouflaged and not startle game, as with retriever breeds, a bland or solid color that will blend into the surrounding foliage is desired (58,59). Terrier breeds, purposed to run alongside hounds in the pursuit of game or to burrow into the terrain to locate quarry, likewise benefitted from white spotting, allowing both hounds and hunters to more easily distinguish the small dogs from foxes (59). White spotting, resultant of variation at the *MITF* gene (17), is broadly present across all clades. However, reflecting the applicable purpose of the color pattern, the presence of breeds fixed for white markings appears to be greatest in the Pointer/Setter, Spaniel, and Terrier clades (S1 Figure).

Morphologically, preference for a shorter foreface, influenced in part by *BMP3* (29), appears at higher frequency in the toy-breed clades (American Toy, Asian Toy, and Toy Spitz), suggesting a preference for a disproportionately shortened face in smaller companion-purposed breeds. Counter to this is the noteworthy observation of specific European Mastiff and Terrier breeds with near fixation for the *BMP3* variant, potentially presenting an argument for production of an intimidating countenance and/or improved jaw strength, as would be appealing for the historic application of those breeds.

### Geography and lineage influence allele distribution among same-breed populations

Dog breeds, while standardized and often interchangeable around the world, have previously been shown to differ in their genetic composition based on geographic locale (34,62–65). We detected significant (p ≤ 0.00167) allele distribution differences based on geographic site of sample collection in 17 of the 26 breeds for which genomic sub-populations could be identified based on geography. Likewise, six breeds (BEAG, ESSP, ECKR, GREY, LAB, WHIP) encompassed multiple lineages differentiated by the application of the dogs to hunting, racing, or conformation competitions. In each case, at least one of the genes queried showed significant allele distribution differences between lineages. These disparate allele frequencies can represent either regional differences in preference or influence of prominent ancestor bias.

*TYRP1* is a representative example. The various recessive alleles of *TYRP1* all ultimately produce the same pigmentation shade (brown instead of black), regardless of the specific alleles present. Therefore, significant variation between *TYRP1* recessive allele frequencies relative to population may indicate a founder genotype or the effect of an influential ancestor. The frequency of the *b^c^* allele is significantly higher in US Field Beagles (p ≤ 0.003) and UK English Springer Spaniels (p ≤ 0.003), compared to conformation lineage Beagles and US English Springer Spaniels which each have higher frequencies of *b^s^*. Conversely, the *b^s^* allele is at a higher frequency in UK conformation lineage Labrador Retrievers (p ≤ 0.003) compared to field-style and US conformation lineage Labrador Retrievers, and UK German Shorthaired Pointers (p ≤ 0.00167) compared to US German Shorthaired Pointers (S2 Figure). As the specific *TYRP1* allele cannot be purposefully selected for through phenotype observation alone, these differences between populations may reflect divergent selection of the lineages. There is no apparent pattern to the distribution of the *b^s^* and *b^c^* alleles relative to clade distinctions (S1 Figure). This would imply that these two *b* alleles are substantially old enough so as to have propagated throughout dog types prior to the development of closed-breed populations.

Traditionally, many dog breeds have had their tails surgically docked to a shorter length, predominantly for the purposes of preventing injury to the tail while the dog carries out its defined function. With the changing use of dogs over time, the necessity for tail docking has recently become a source of discussion among canine enthusiasts and the veterinary community. Between 1987 and 2018, 35 countries have banned or restricted tail docking or the ability to exhibit dogs with docked tails in regulated kennel club events (66–70). There are currently no restrictions placed on docking in the US, while the UK has banned tail docking since 2006 (66). Dog breeders in countries with docking bans wanting to uphold the traditional appearance of a dock-tailed breed could therefore select for the *tailless* variant of the *T* gene to produce naturally shortened tails in their dogs. However, none of the breeds from the US and the UK that were genotyped in the present study show significant differences in the *tailless* variant frequencies. Eleven breeds collected in both locations traditionally have docked tails, while only four of these breeds (PEMB, SKIP, VIZS, WELT) had any measureable level of the *tailless* variant present. While the level of *tailless* is numerically higher in the UK populations of the Pembroke Welsh Corgi, Schipperke, and Vizsla compared to US populations, the differences were not significantly different (p ≤ 0.00167). This lack of disparity likely reflects the age of the samples used in this dataset, which were collected between 2005 and 2016. It is likely that insufficient time has passed to accurately reflect the efforts of selection for natural taillessness resulting from the procedural ban, and the *T* allele frequencies are expected to increase in select breeds over time in those countries banning surgical docking.

It is worth noting that, while overall numbers for most breeds were quite high, once split into geographic or lineage groups, the cohort size in some cases became smaller. For example, this study includes 111 Beagles, but when split into US Show Beagles (n = 43), US Field Beagles (n = 52) and UK Beagles (n = 16), the UK population of Beagles was relatively small. Another way to consider this data is by ratios; for example, the 380 Labrador Retrievers are split into three populations with a ratio of approximately 1:2:1, while the 50 Schipperkes are divided into two populations with a ratio of approximately 1:9. In only two instances, however, does a sample sub-population represent less than 10% of the total breed sample population size (UK Soft Coated Wheaten Terriers represent 9.6% of the total for the breed, Medium Poodles represent 6.4% of the total for that breed). These small numbers may ultimately influence the accuracy of allele frequency estimates in these subpopulations, and larger sample sizes in future studies would solidify true differences between such populations.

### The natural probability of disallowed traits in pure breeds

A breed standard is a written description of a given breed, as determined by a committee of educated breeders, that details how a typical dog of that breed should look and behave. Historically, the standards functioned as a framework for breeders to aim at when producing dogs of that breed. Breed standards outline traits that are disallowed, so that breeders can opt to breed away from or minimize their occurrence. Each canine registering organization, usually specific to country or region, employs its own set of breed standards and, for the most part, these standards are concordant breed-to-breed across registries. However, there are instances where the wording between standards is not precisely mirrored, such that variations in color, size, or morphology are more or less tolerated from registry to registry.

Our analyses detected genetic variants that would cause disallowed phenotypes in 67.5% of breeds tested (S3 Table). The majority of these, 58.9%, have a <1% probability of producing that undesirable phenotype given random breeding. These values represent multi-generational efforts to eliminate unfavorable phenotypes within breeds, however they also highlight that conformity to breed standard is certainly not yet universal or complete. They also illustrate the difficulty in selecting against alleles that are masked not just by dominance, but also epistasis, even in a highly-visible trait such as coat color. In general, these findings exemplify three separate scenarios: 1) traits broadly disallowed but carried at low frequencies, 2) traits allowed under some registries but not others, and 3) single traits that persist due to breed-specific allowances for particular trait combinations (Figure 5). For example, regardless of breed registry (AKC, UKC, KC, and FCI), the Bull Terrier is described as always having a black colored nose, and never showing brown hair pigmentation. However, we detected the *b^c^* and *b^s^* alleles of *TYRP1* at a frequency of 3% each in our population of pure bred Bull Terriers. This results in a probability of 0.0036 in producing a brown colored purebred Bull Terrier, present as brown pigmented skin and/or coat, assuming random breeding (Figure 5; S3 Table). Conversely, Shetland Sheepdog breed standards differ in that the AKC disallows piebald spotting in the breed, the FCI and KC tolerate, but do not prefer, excessive white spotting, and the UKC allows any variation from no white to fully piebald. We detected the *MITF* variant for white spotting, known to cause the piebald phenotype in this particular breed (18), at a frequency of 16% in the UK population of the breed and 6% in the US population (Figure 5; S2 Table and S3 Table). Finally, the Great Dane breed standard is relatively cohesive across breed registries, but defines acceptable coat colors in terms of pattern combinations; for example, white spotting and the harlequin pattern are only allowed on a black base color. However, because black base color is dictated by *CBD103* on chromosome 16 with a breed frequency of 66%, white spotting is controlled by *MITF* on chromosome 20 with a breed frequency of 6%, and harlequin is caused by *PSMB7* on chromosome 9 with a breed frequency of 21%, the realistic potential to produce a fawn-based harlequin or a fawn and white spotted Great Dane is unavoidable (Figure 5, S2 Table). These values result in a 1.33% probability of producing a fawn and white Great Dane through random breeding of purebred dogs, or a 3.80% probability of producing a fawn Great Dane with one copy of the harlequin mutation. Since the frequency levels of the merle variant – required for production of the harlequin phenotype – are not known, the latter value does not necessarily reflect the number of fawn-based harlequin dogs produced.

**Figure 5:**
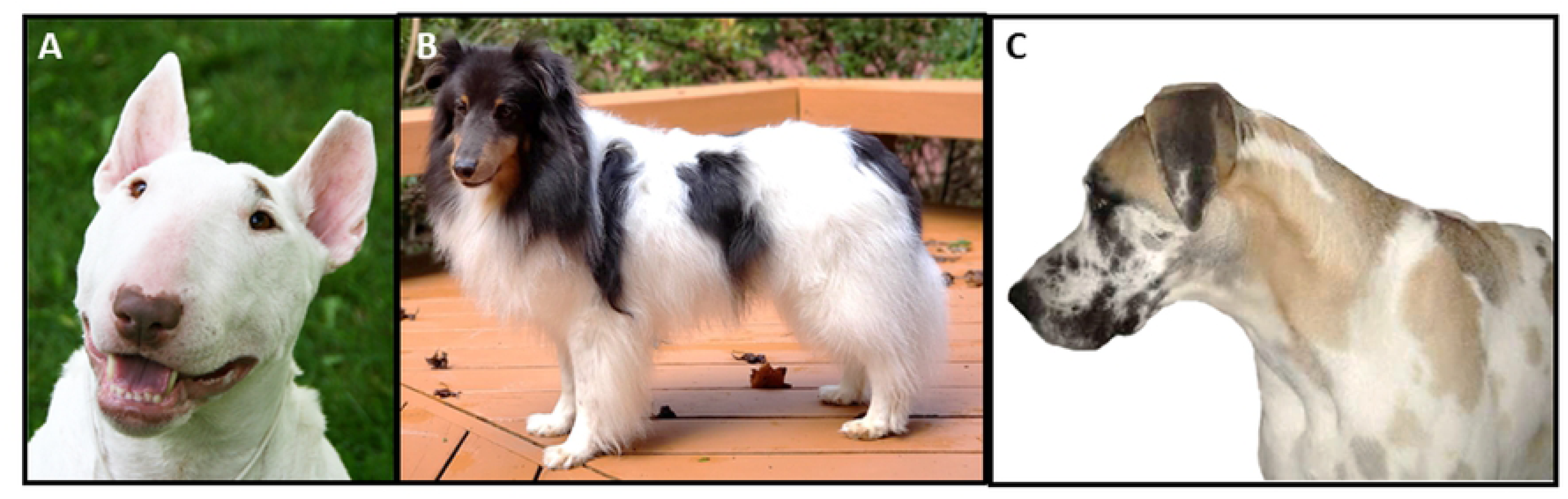
Purebred dogs exhibiting color traits deemed inappropriate by one or more breed registries. A) Bull Terrier with a brown nose and brown patch above its eye. B) Shetland Sheepdog with piebald white spotting. C) Great Dane with the harlequin pattern on a fawn base color.

The Schipperke breed, collected from populations in the US and the UK, presents a clear example of how regional acceptance of certain characteristics can drive the frequency of a variant within a population. The Schipperke breed standards for AKC, UKC, and FCI all state that the dog must be entirely black in color. However, the KC in the UK states that “any solid color” is permissible. As such, the allele frequencies for the *MC1R* recessive *e* allele, which produces a solid red color when homozygous, was observed at 50% among the dogs sampled in the UK (n = 6), and 0% among the dogs sampled in the US (n = 44) (Figure 2).

Much recent emphasis has been placed on the importance of genetic diversity within breeds (71–78). With the conservation of diversity in mind, breeders and breed organizations must weigh the relative value of breed standard conformity with preservation of genetic diversity. The existence of unfavorable, though arguably benign pigmentation or morphological variations, has here been quantified and can be addressed by applied genetic screening to reduce the carrier frequency of breeding stock, or by reassessing breed standards to broaden the acceptance of preexisting variation. Likewise, though our analyses have indicated that production of disallowed phenotypes is generally quite low, the occurrence of an undesirable pigmentation trait should not necessarily exclude a dog from purebred status if that variant has been detected in the appropriate population.

### Diverse breed representation of rare alleles informed by ancestral haplotype sharing

The unique structure of the dog genome dictates that, even across breeds, large regions of linkage disequilibrium can accompany trait-causing variants (79), as measured by IBD haplotype sharing. As such, we can assume that the presence of a given allele in two breeds may indicate shared ancestry between those breeds. For example, it has long been assumed that the recessive black allele of *ASIP* (*a*) is predominantly found in herding breeds (3–5). However, among the 212 breeds analyzed here, the *a* allele was identified in 89 breeds, 83 of which are not previously reported as carrying the allele, representing 23 of the possible 28 clades assigned using IBD haplotype sharing in previous work (34,35). While herding breeds, present in the Continental Shepherd, Hungarian, New World, Nordic Spitz, and UK Rural clades, comprise 14 of the *a* allele-possessing breeds – an expected result – the greatest breed representation was among the Pointer/Setter clade with 13 breeds possessing the *a* allele. The only clades without measured frequency of the *a* allele are the Alpine, American Terrier, Asian Toy, Pinscher, and Standard/Miniature Schnauzer clades.

IBD haplotype sharing can reflect a shared ancestry between populations, and the amount of haplotype sharing between breeds correlates significantly with the time point at which those breeds shared a common ancestry (34). Using instances of significant levels of haplotype sharing between breeds, with reliably dating introgression events to as early as the late 1800’s (34), 65 of the 89 breeds displaying carrier frequencies >0% for the *a* allele can be connected (Figure 4). While not necessarily reflecting the exact mode of allele sharing between breeds, the measured haplotype sharing instances successfully demonstrate a recent ancestral history between the breeds. There are 21 breeds with positive *a* allele frequencies that cannot be connected via haplotype sharing due to not being included in the previous haplotype sharing analyses (34), but are predicted to be in phylogenetic clades already represented among *a*-carrying breeds. Three breeds, the Anatolian Shepherd, Kuvasz, and Tibetan Mastiff, carry the *a* allele and are included in the haplotype sharing analyses, but do not show significant ancestry with other breeds. Therefore, while the extensive 65-breed relationship matrix supports the potential for the *a* allele to spread between breeds via recent introgression events, the presence of the allele in the Anatolian Shepherd, Kuvasz, and Tibetan Mastiff, for which no recent introgression events have been detected, suggests that the allele itself arose early in the history of the domestic dog, establishing a broad distribution well before the development of modern breeds in the late 1800’s. Support for the persistence of the *a* allele in the observed 89 breeds lies in recognizing that at least 55 of the breeds prefer a solid eumelanin (black, brown, or grey) base color. Selection for solid eumelanin would favor both the *K^B^* allele of *CBD103* and the *a* allele of *ASIP*. A further 18 of the observed breeds display phenotypes that are incompatible with the *a*/*a* recessive black phenotype, allowing it to remain mostly undetected: 11 of the breeds are solid white (AESK, BBLS, BICH, BOLO, COTO, JSPZ, KUVZ, MALT, MARM, SAMO, VPIN), five breeds are fixed at *MC1R e*/*e* red (GOLD, ISET, SPIN, VIZS, WVIZ), and two breeds display an extreme piebald white spotting pattern with minimal concern for the residual base color (DALM, GPYR). The remaining 16 breeds have an *a* allele frequency of < 10%, allowing for the production of the recessive trait at rates deemed inconsequential to influence selection decisions by breeders.

The *tailless* allele of the *T* gene was identified in 48 breeds, representing 14 clades. While 10 of these breeds have been previously identified as carriers of the *tailless* allele (25,26), 38 are reported here for the first time. Thirty-eight of the 48 *T*-carrying breeds are represented in the identity-by-descent dataset (34), all of which can be connected into a single relationship matrix (Figure 3). In some instances, such as with the Brittany and Newfoundland, which have *tailless* carrier frequencies of 0.04 and 0.01 respectively, there is no direct identity-by-descent relationship with a potential source breed of taillessness. However, both show ancestral relationships with the Golden Retriever, who then further shows ancestry with the Airedale Terrier, a carrier of *tailless* at a frequency of 0.02. It is possible that the Golden Retriever either carries *tailless* at a frequency not detected in our screening, or the trait previously existed within the breed and has since been selected against and eliminated. Other possible historic sources of *tailless* that have decreased or eliminated their current carrier frequency include the Icelandic Sheepdog, Keeshond, Mastiff, and Pug.

Variation at *FGF5*, responsible for the long and short haired phenotypes in Pembroke Welsh Corgis, Border Collies, Dachshunds, and German Shepherd Dogs (80), presented across our 212 breeds primarily as expected. Thirty-eight breeds had 100% frequency of the longhaired variant, together with phenotypes ranging from body-wide presence of hairs with length sufficient to reach the ground (eg. Komondor, Havanese) to long hair production predominantly on the ventral surfaces of the limbs, tail, and body (eg. Brittany, Borzoi). Thirty-five breeds had 100% frequency of the shorthaired variant, including breeds with notably the shortest hair phenotypes (eg. Boxer, Doberman Pinscher), but also including many breeds with traditionally longer, often wire-textured, hair on their limbs and muzzles (eg. Berger Picard, Otterhound, Lakeland Terrier), or plush double coats (eg. Norwegian Elkhound, Norwegian Buhund). Indeed, some breeds known for their abundant coats, such as the Afghan Hound, Tibetan Mastiff, or Chow Chow, expressed the *FGF5* longhaired variant at the relatively low frequencies of 21%, 49%, and 61%, respectively. Similar outcomes were observed with regard to the curly coat variant of *KRT71*. Eight breeds showed 100% fixation for the curly variant, including the Irish Water Spaniel, Komondor, and Airedale Terrier. Forty-nine breeds genotyped at 100% frequency for wild-type (no curl) *KRT71*, ranging from the long, straight-haired Gordon Setter, to the smooth-coated Miniature Pinscher. Of great interest is the fixation of the wild-type (no curl) variant in breeds such as the Curly-Coated Retriever, Bouvier des Flandres, and Otterhound, which traditionally have coats with noticeable curl. In some instances, appreciable levels of the curly coated variant were detected in breeds not expected to have curly hair, such as the German Pinscher (24%), Smooth Fox Terrier (16%), or Catahoula Leopard Dog (12%). In these instances, the breeds generally have insufficient hair length to present any amount of curl, allowing the variant to persist undetected. While the measured variants in *KRT71* and *FGF5* account for some of the variation in coat type, these newly reported allele frequencies highlight the genetic complexity involved in hair production. Indeed, recent research has identified additional variants, not covered in the scope of this project, that account for coat variation previously unexplained by the original *KRT71* and *FGF5* variants (22–24). Clearly, extreme coat type phenotypes exist that are outside of the control of the known variants, two of which were explored here.

Lastly, while the allele frequencies reported herein are intended to provide general information regarding the existence of particular alleles within breeds, they are not likely to be perfectly accurate estimates of actual breed-wide allele frequencies. For instance, while the German Shepherd Dog, Lagotto Romagnolo, and Australian Shepherd are represented by 162, 139, and 137 dogs, respectively, our dataset only includes eight Small Munsterlanders, and nine each of Redbone Coonhounds and Bergamascos. As mentioned above, this is likewise true for the regional or functional same-breed populations. Despite these limitations, no other published genotype dataset matches the presently reported size and diversity of dog samples across a dozen pigmentation and morphologically relevant genes.

### Conclusion

The broad adoption of commercial genotyping services yields an immense amount of genetic information. We have demonstrated how this data can be utilized to detail the current phenotypic diversity of 212 dog breeds, and the impact of population divergence due to geographic separation or selection practices. While most dog breeds have existed as closed breeding populations since the late 19^th^ century, we have shown that rare trait-causing variants continue to persist within most breeds. Conflicting breed standard descriptions of multiple registering organizations may have facilitated the persistence of these traits within certain populations. In addition, epistatic masking effects have contributed to the continuance of various trait-causing alleles, due to the complicated multi-gene pathways whereby recessive alleles can remain unexpressed over multiple generations, and making selection for or against them challenging without the use of genotyping services.

The modern existence of domestic dogs is such that coat color is primarily a matter of aesthetics. Consideration should be given by breed associations to unifying breed standards across registering bodies, either with the intent of increasing selective pressure against truly undesirable characteristics, or expanding the standards to permit phenotypes for which the causal variants exist ancestrally within the breed. Simultaneously, conservation of genetic diversity within breeds must be weighed. The present study documents for the first time the frequencies of alleles at 12 coat color and morphological genes, across 11,790 dogs, representing 212 breeds, and demonstrates not only the anticipated genotypic variations within breeds, but also rare and unexpected alleles not previously reported.

## Materials and Methods

### Sample collections

Genotype data from 11,790 canids, representing 212 pure dog breeds and 4 wild canine populations, was compiled during the development and implementation of the Mars WISDOM PANEL platform (Wisdom Health, Vancouver, WA, USA). DNA collections herein represent a subset of those initially reported in Donner, *et al.* (43). Dog DNA samples were obtained by Wisdom Health (formerly Mars Veterinary) and Genoscoper Laboratories (Helsinki, Finland), between January 2005 and October 2016, as owner-submitted, non-invasive cheek swab collections. Dog owners provided consent for use of their dog’s DNA in research. The dogs sampled predominantly originated from the US and UK, though samples were also obtained in smaller numbers from several other countries. Dogs were considered to be purebred if registered with a relevant all-breed registry, the predominant ones being: Fédération Cynologique Internationale, American Kennel Club, United Kennel Club, and UK Kennel Club, or an applicable single-breed registry for rare breeds. Breed and species classification was further verified through principal component analysis (PCA) and genotyping on the WISDOM PANEL platform (Wisdom Health), particularly for the minority newer/rarer breeds not currently (at the time, or even now) recognized by a national registry. PCA further revealed that 30 breeds formed sub-populations based on geography, body size, coat type, or function (e.g., show vs. field). Archived wild canine samples were used to represent: 1) the grey wolf (*Canis lupus lupus*, sampled primarily from Eastern North America, n = 12); 2) coyote (*Canis latrans*) collected from Eastern North America (Eastern coyote, n =29) or British Columbia and Southwestern US (Western coyote, n = 19), populations segregated as determined by PCA cluster analysis; and 3) dingo (*Canis lupus dingo*) (n = 12) populations. All dog breeds were represented by ≥10 individuals, with the exception of the Small Munsterlander (n = 8), Bergamasco (n = 9), and Redbone Coonhound (n = 9).

### Genotyping

Genotyping of seven coat color and five morphological trait variants (Table 1) was conducted on a custom-designed Illumina Infinium HD bead chip using manufacturer-recommended protocols ((81); Illumina, San Diego, CA, USA). The validation and genotyping quality control measures for this platform were previously described in detail (43,81). Trait variant assays specifically were valideated through extensive correlation of genotypes with established breed phenotypes and owner-submitted pictures of individual dogs.

### Allele Frequencies and Statistical Analysis

Allele frequencies for each variant were determined for each breed and, when appropriate, breed subpopulations, and converted to binary genotypes for genes with multiple alleles. The total number of dog samples per breed are reported in S1 Table. Due to sporadic failure, the number of genotypes obtained per breed for each gene may vary from the total number of dogs per breed (S2 Table). For breeds with subpopulations (n = 30 breeds, delineated primarily by geography, but also body size, breeding line of conformation versus working, etc.), within-breed statistical significance of the differences in genotype distribution between the subpopulations was evaluated by Pearson’s chi-square contingency tables, or Fisher’s exact tests (when allele counts in any single cell fell below 5), for each gene. Calculations were conducted with the MASS package in R (82,83). A p-value of < 0.00167 was chosen to indicate a significantly different allele distribution between the same-breed subpopulations; this is the Bonferroni correction for multiple testing of 30 breeds for each gene (i.e., 0.05/30). In the case of *FGF5*, no genotypes were available for the US population of German Spitz, thus significance was corrected instead for 29 breeds, to a p-value of ≤ 0.00142 for this gene. Further correction (e.g., for testing of 12 different genes) was not applied because a balanced approach was desired and the initial Bonferroni correction was deemed suitable. Further analysis was conducted for the six breeds sampled from > 2 subpopulations. When these breeds differed significantly at any gene using the initial analysis described above, they were subsequently evaluated for pairwise significance using Pearson’s chi-square or Fisher’s exact tests. Significance for each breed subpopulation was determined by p-value = 0.05/n, where n = maximum number of subpopulations remaining in analysis. This number therefore varied for each gene, depending on remaining significant breeds. For example, if Poodles – divided into four subpopulations – remained, the correction was 0.05/4, resulting in a p-value cut-off of 0.0125; however if Dachshunds – divided into six subpopulations – remained, the correction was 0.05/6, resulting in a p-value cut-off of 0.0083. Specific statistical applications and n values are indicated in the appropriate figure, table legends or footnotes.

### Probability of phenotype expression

When alleles that would produce an undesirable or disallowed phenotype were observed in a given breed, relative to AKC, UKC, KC (Britain), or FCI breed standard descriptions, the probability of producing that phenotype was calculated with the following equation:

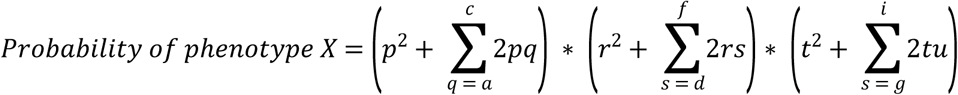

Such that:

*p* = the frequency of the fault-producing allele, *P*, of the gene causal for *X*

*q* = the frequency of any same-gene allele recessive to *P*, of which there can be up to 3 (*a* to *c*)

*r* = the frequency of the most dominant allele, *R*, at an interacting gene that is required for production of the fault phenotype

*s* = the frequency of any same-gene allele recessive to *R*, of which there can be up to 3 (*d* to *f*)

*t* = the frequency of the most dominant allele, T, at an interacting gene that is required for production of the fault phenotype

*u* = the frequency of any same-gene allele recessive to T, of which there can be up to 3 (g to i)

The traits for which genotypes were obtained present scenarios in which up to two known interacting genes can influence the expression of a phenotype. Namely, expression of an *ASIP* phenotype relies on corresponding *CBD103* and *MC1R* genotypes, the *MC1R* phenotypes caused by *E^M^* and *E^G^* require specific genotypes at *ASIP* as well as a homozygous wild-type *CBD103* genotype, and a recessive genotype at *CBD103* can result in multiple possible *ASIP* phenotypes, only some of which may be undesirable. Conversely, recessive homozygosity at *TYRP1* will result in brown pigment of the hair and keratinized skin, which may present as a coat fault if coupled with eumelanin-producing *MC1R* and *ASIP* genotypes, or a nose pigment fault regardless of hair pigmentation. Taillessness is only expressed in the heterozygous state, as it is embryonic lethal when homozygous, therefore, probability values were corrected to reflect outcomes among live births.

## Acknowledgements

The authors gratefully thank all the dog owners, breeders, and veterinarians who contributed to this study by testing their dogs and through their interest in advancing canine genetics research. They would also like to thank Dr. Hsin-Yie Weng for statistical assistance and Stephen Davison for classification of sub-populations.

**Supporting information captions** (files uploaded separately)

**S1 Figure:** Distribution of alleles for a) *TYRP1*, b) *MITF*, c) *PSMB7*, d) *RALY*, e) *KRT71*, f) *FGF5*, g) *T*, h) *BMP3*, i) chr10 ear set marker. Breeds are grouped by phylogenetic relationship.

**S2 Figure:** Allele frequencies for a) *MC1R*, b) *ASIP*, c) *TYRP1*, d) *CBD103*, e) *MITF*, f) *PSMB7*, g) *RALY*, h) *KRT71*, i) *FGF5*, j) *T*, k) *BMP3*, l) chr10 ear marker, for all breeds with multiple populations. Initial X^2^ significance (p < 0.0167) is indicated by horizontal black bars. Pairwise significance was conducted for all significant breeds with greater than two populations, with significance level indicated by **. Letters under horizontal black bar denote significant groupings.

**S1 Table:** List of breeds, their abbreviations used throughout the paper, and the number of samples genotyped. Each breed was assigned to a phylogenetic clade based on previously published results (34,35). Clade names in parenthesis indicate breeds not included previously (34,35), but for which a clade was assigned based on known breed history and phenotypes.

**S2a Table:** Allele frequencies for coat color genes *ASIP* and *MC1R*. Breeds fixed for a single allele at any gene are indicated with bold text.

**S2b Table:** Allele frequencies for the brown (*TYRP1*) and dominant black (*CBD103*) genes. Breeds fixed for a single allele at any gene are indicated with bold text.

**S2c Table:** Allele frequencies for the white spotting (*MITF*), harlequin (*PSMB71*), and saddle tan (*RALY*) genes. Breeds fixed for a single allele at any gene are indicated with bold text.

**S2d Table:** Allele frequencies for hair length (*FGF5*), hair curl (*KRT71*), and ear set. Breeds fixed for a single allele at any gene are indicated with bold text.

**S2e Table:** Allele frequencies for skull shape (*BMP3*) and natural taillessness (*T*). Breeds fixed for a single allele at any gene are indicated with bold text.

**S3 Table:** Breeds genotyped to have alleles that would produce phenotypes considered as a “fault” by either the American Kennel Club (AKC), Fédération Cynologique Internationale (FCI), United Kennel Club (UKC), or The Kennel Club of the UK (KC). The level of tolerance within each breed registry is designated as either not allowed (N), not preferred (n.p.), allowed (Y), or ambiguously worded (amb.). A breed not recognized by a given organization is indicated with a dash (-). Inheritance of the fault-causing allele is designated as dominant (D), recessive (R), or compound heterozygote (CH). Breed name abbreviations are as listed in S1 Table. Probabilities for producing the non-standard phenotype were calculated assuming random mating within the breed, and account for multi-gene inheritance, expression, and epistatic effects.

